# Structure of the archaellar motor and associated cytoplasmic cone in *Thermococcus kodakaraensis*

**DOI:** 10.1101/108209

**Authors:** Ariane Briegel, Catherine M. Oikonomou, Yi-Wei Chang, Andreas Kjaer, Audrey N. Huang, Ki Woo Kim, Debnath Ghosal, Robert P. Gunsalus, Grant J. Jensen

## Abstract

Archaeal swimming motility is driven by rotary motors called archaella. The structure of these motors, and particularly how they are anchored in the absence of a peptidoglycan cell wall, is unknown. Here, we use electron cryotomography to visualize the archaellar motor *in vivo* in *Thermococcus kodakaraensis*. Compared to the homologous bacterial type IV pilus (T4P), we observe structural similarities as well as several unique features. While the position of the cytoplasmic ATPase appears conserved, it is not braced by linkages that extend upward through the cell envelope as in the T4P, but rather by cytoplasmic components that attach it to a large conical frustum up to 500 nm in diameter at its base. In addition to anchoring the lophotrichous bundle of archaella, the conical frustum associates with chemosensory arrays and ribosome-excluding material and may function as a polar organizing center for the coccoid cells.

## INTRODUCTION

Motility is a fundamental property of single-celled organisms. In archaea, swimming motility is driven by a rotary motor called the archaellum. Archaella are functionally analogous to bacterial flagella, but evolutionarily homologous to the type IV pilus (T4P) and type II secretion system (T2SS) machineries of bacteria [1]. Recently, an atomic structure of the archaellum fiber purified from the euryarchaeon *Methanospirillum hungatei* revealed differences compared to the bacterial T4P fiber, including lack of a central pore and more extensive inter-subunit interactions [2]. The structure of the archaellar basal body, and its similarity to the T4P basal body remains unknown.

Unlike T4P fibers that only assemble and disassemble, archaella assemble and can then rotate in both directions to either push or pull the cell [3, 4]. Light microscopy of *Halobacterium salinarum* revealed discrete steps during rotation, likely corresponding to ATP hydrolysis events by the basal body ATPase, FlaI [5]. While the bacterial T4P contains two distinct ATPases for assembly and disassembly of the pilus fiber, the single ATPase FlaI drives both assembly and rotation of the archaellum [6]. The N-terminal domain of the archaellum/T2SS/T4P superfamily ATPases is the most variable, and the first 29 residues of FlaI, located on the outer edge of the hexamer, were found to be essential for motility but not assembly, although the basis of this functional separation remains unclear [6].

FlaI is predicted to interact with the integral membrane protein FlaJ [7]. Structural studies of the bacterial T4P suggest that ATPase-driven rotation of the FlaJ homolog, PilC, incorporates pilin subunits from the membrane into the growing fiber [8]. This is possible because the ATPase itself is clamped in an integrated structure that spans the inner and outer membranes and periplasm and anchors on the cell-encompassing peptidoglycan cell wall [8]. A similar cell-wall-attached structure anchors the rotation of the bacterial flagellar motor [9]. Without knowing the structure of the archaellar basal body, it is unclear how similar anchoring could occur in the envelope of archaea, which consists of a single membrane and thin proteinaceous surface (S-)layer. It was recently proposed that FlaF might anchor the archaellum through interactions with the S-layer [10]. Others have suggested that a cytoplasmic structure mechanically stabilizes the motor [3]. Supporting this idea, cytoplasmic structures underlying the archaella have been observed by traditional electron microscopy (EM) of Halobacteria [11, 12].

Electron cryotomography (ECT) can image intact cells in a frozen, fully-hydrated state, providing macromolecular-resolution (~4-6 nm) details about native cellular structures [13]. Here, we used ECT to visualize the structure of the archaellar basal body *in vivo* in *Thermococcus kodakaraensis* cells. *T. kodakaraensis* (originally designated *Pyrococcus sp*. strain KOD1 and later identified as belonging to the *Thermococcus* genus [14]; also known as *T. kodakarensis*) is one of the best-studied archaeal species. It was isolated from a Japanese solfatara in 1994 [15], and has proven readily amenable to genetics (well-developed gene manipulation techniques exploit its natural competence [16]) and the isolation of thermostable enzymes (e.g. high-fidelity DNA polymerase for PCR [17]). In addition to revealing the overall structure of the archaellar basal body *in vivo*, we discovered a novel cytoplasmic conical structure in *T. kodakaraensis* associated with archaellar motility and potentially other polar organizing activities.

## RESULTS

We imaged *T. kodakaraensis* cells by ECT in a native, frozen-hydrated state. Many cells appeared to be lysed prior to plunge-freezing for ECT, but out of 18 apparently intact cells, we observed a lophotrichous bundle of archaella in 13. Each bundle contained between four and 14 archaella. Due to the relatively large size of *T. kodakaraensis* cells (cells are irregular cocci ~1.5 µm in diameter), only a portion of the cell was visible in the limited field of view of our high-magnification cryotomograms. We therefore think it likely that in the remaining five cells, the archaellar bundle was present but not located in the portion of the cell imaged. In addition, we observed well-preserved archaellar bundles in eight apparently lysed cells.

We consistently observed a prominent conical structure associated with the archaellar bundle in the cytoplasm (Figure 1). We never observed archaella unassociated with a cone, or vice versa. The conical structure showed a consistent morphology and localization inside the cell: closely associated with, but not touching, the cytoplasmic membrane at its narrow end and expanding a variable length to a wide base, which varied from 220 to 525 nm in diameter. The central axis was perpendicular to the membrane, as seen in cross-sectional side views (Figure 1A–C, additional examples in Figure EV1). The edges, seen in cross-section, frequently exhibited periodic densities suggestive of individual protein subunits (Figure 1C), with a thickness of 3-4 nm. We observed that while cones in different cells had different heights, the opposite edges within each cone were always symmetric (of similar lengths). Tomographic slices capturing the central axis of the conical structure in side view showed an angle of 109 ± 6^o^ (mean ± s.d., n=5) between opposite edges. The structures were not complete cones but rather conical frusta: they did not taper fully to a point, but exhibited a blunt tip. In top-views, we observed a ring situated in the throat of the frustum, just below the tip (Figure 1D, E). These rings comprised 19 subunits (Figure 1D inset), each again 3-4 nm thick, with an overall ring diameter of 31 ± 2 nm (mean ± s.d., n=10). The position of the ring in the conical frustum was clearest in tomograms of lysed cells, which were thinner and contained less cytoplasmic material (Figure 1F–H). Even in such tomograms, however, we could not visualize a well-defined connection between the two portions of the structure, so it is unclear if and how the components are connected.

**Figure 1.**
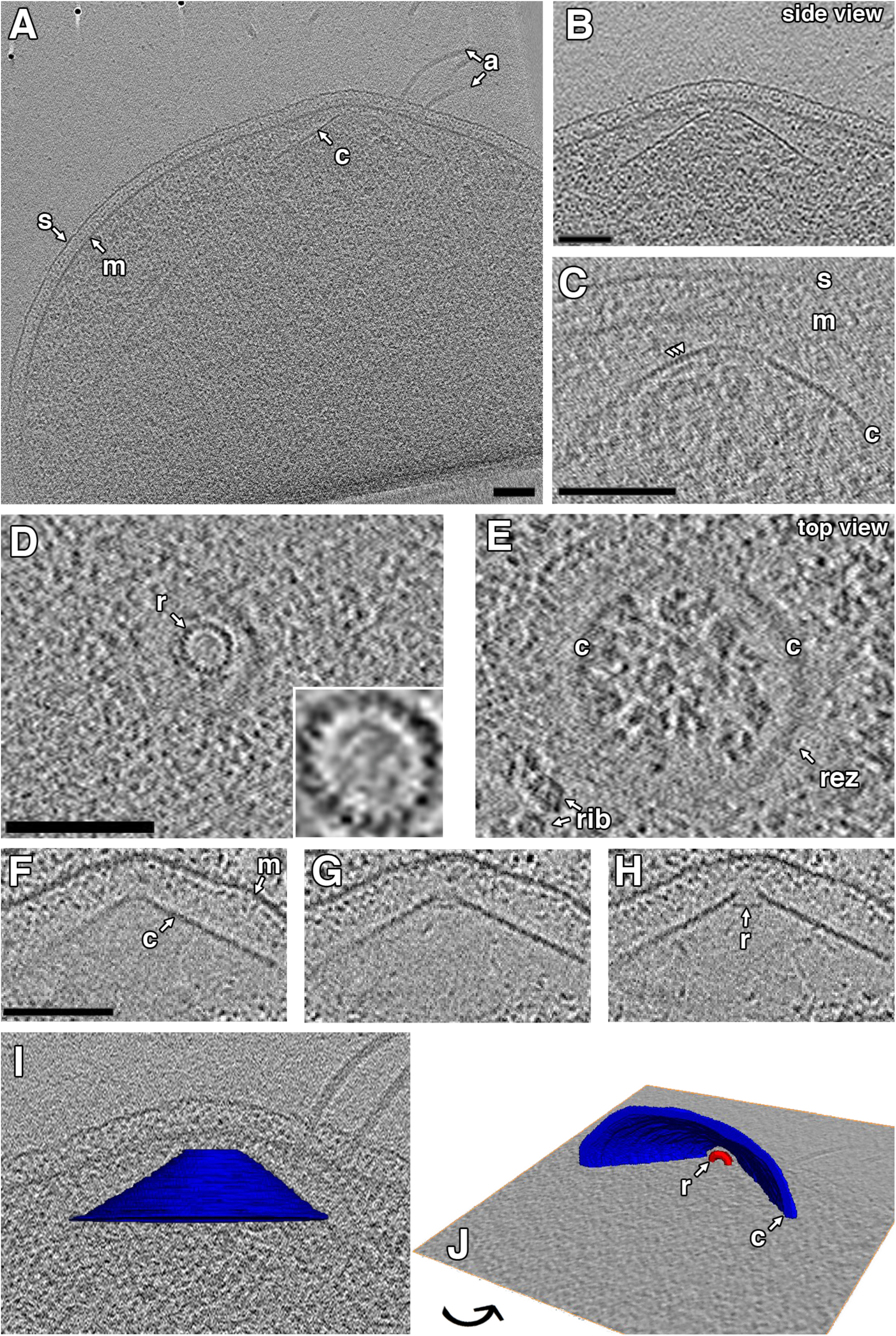
Cytoplasmic conical structures in *Thermococcus kodakaraensis*. **(A)** Atomographic slice shows a side view of a conical structure (c) in the cytoplasm, rotated and enlarged in **(B)**. **(C)** A tomographic slice shows a side view of the cone in another cell, highlighting the subunit texture along the edge of the cone (arrowheads). **(D, E)** Top views of a cone at different heights show the inner ring (r; enlarged in inset to highlight 19-subunit structure) and outer cone. **(F-H)** Sequential slices through a side view of a cone in a lysed cell show the relative location of the ring in the cone. **(I, J)** Different views of a 3D segmentation of the cone shown in **(A)**, embedded in a tomographic slice. s, S-layer; m, membrane; a, archaella; rib, ribosomes; rez, ribosome-excluding zone. Scale bars 100 nm; segmentation not to scale.

Conical structures were surrounded by an ~30-45 nm wide ribosome-excluding zone (REZ) (Figure 1E, Figure 2). In nearly all cells, both intact and lysed, we observed filament bundles near or associated with this REZ (Figure 2B-E). The bundles were more extensive in lysed cells. Each filament was ~12 nm wide and made up of a series of disk-like densities spaced ~7 nm apart. Chemosensory arrays were also consistently observed near the conical structures (Figure 2A, B). In one cell, we observed two attached conical structures, each associated with archaella and each approximately 250 nm in diameter at its base (Figure EV2).

**Figure 2.**
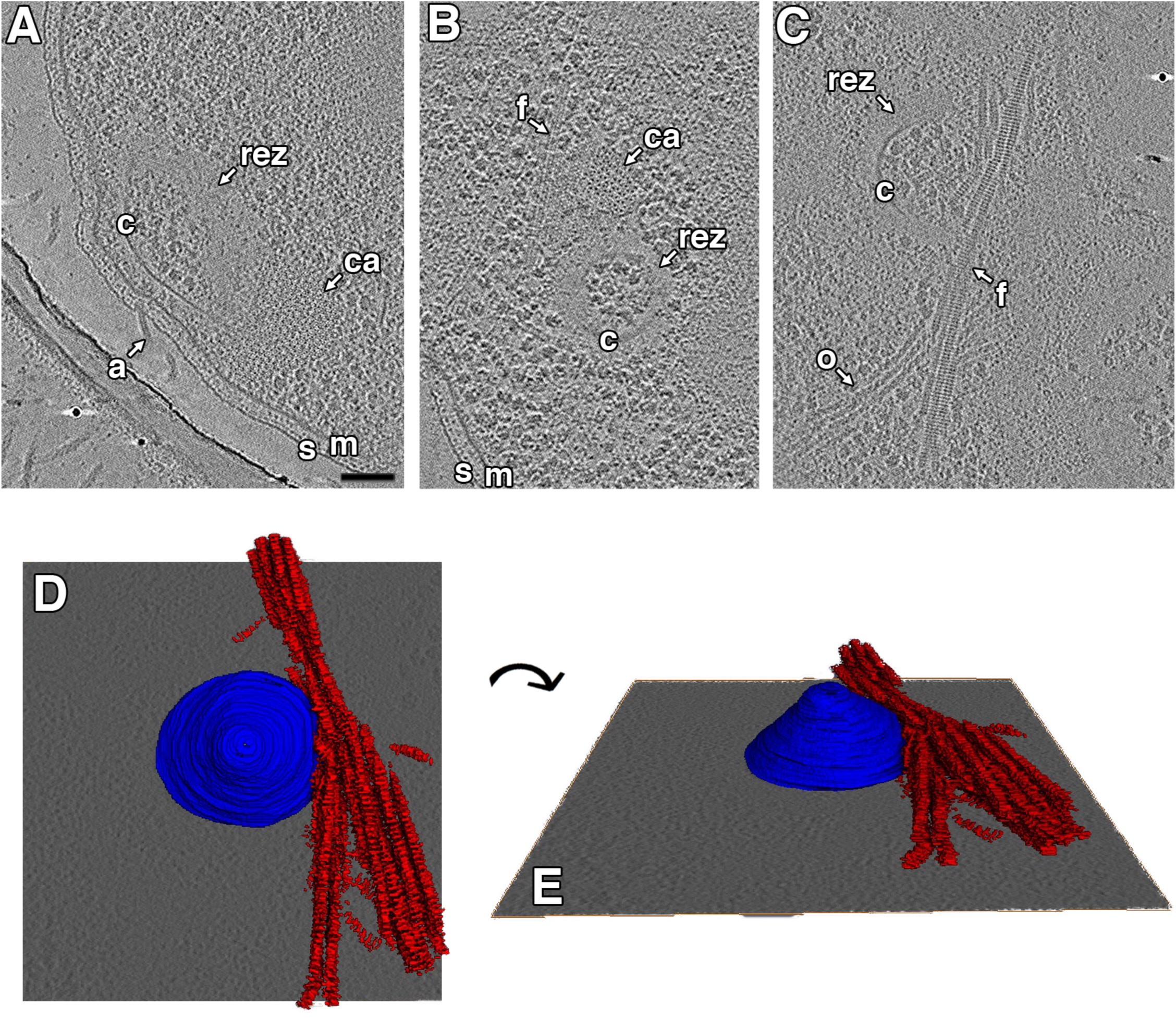
Cones are associated with chemosensory arrays, ribosome-excluding zones and filament bundles. Tomographic slices show side **(A)** and top **(B, C)** views of cones (c) in threecells, highlighting associated chemosensory arrays (ca), ribosome-excluding zones (rez) and filament bundles (f). For a slice-by-slice view through the tomogram shown in (A), see Movie S1. **(D)** and **(E)** show different views of a 3D segmentation of the structures shown in **(C)**, with the conical structure in blue and the filament bundle in red. s, S-layer; m, membrane; a, archaellum; o, other filaments. Scale bar 100 nm; segmentation not to scale.

To characterize the interaction between the archaellar bundle and the conical structure, we measured the distance from the base of each archaellum in the membrane to the cone. The structure of the cone means that the distance between it and the membrane varies – shortest at the tip of the cone and longest at the base. Since archaella were located at various radial positions along the cone, we expected their distance to vary similarly. Interestingly, however, we measured a much more consistent distance of 44 ± 5 nm (mean ± s.d., n=29) from the cone to the base of each archaellum in the membrane (Figure 3). Consistent with this, we observed a variety of orientations of archaella in the cell envelope, frequently not perpendicular to the S-layer, allowing the conserved distance to the cone (Figure 3, Figure EV3). In a few cases, we observed continuous densities connecting the archaella and the cone (Figure EV3E).

**Figure 3.**
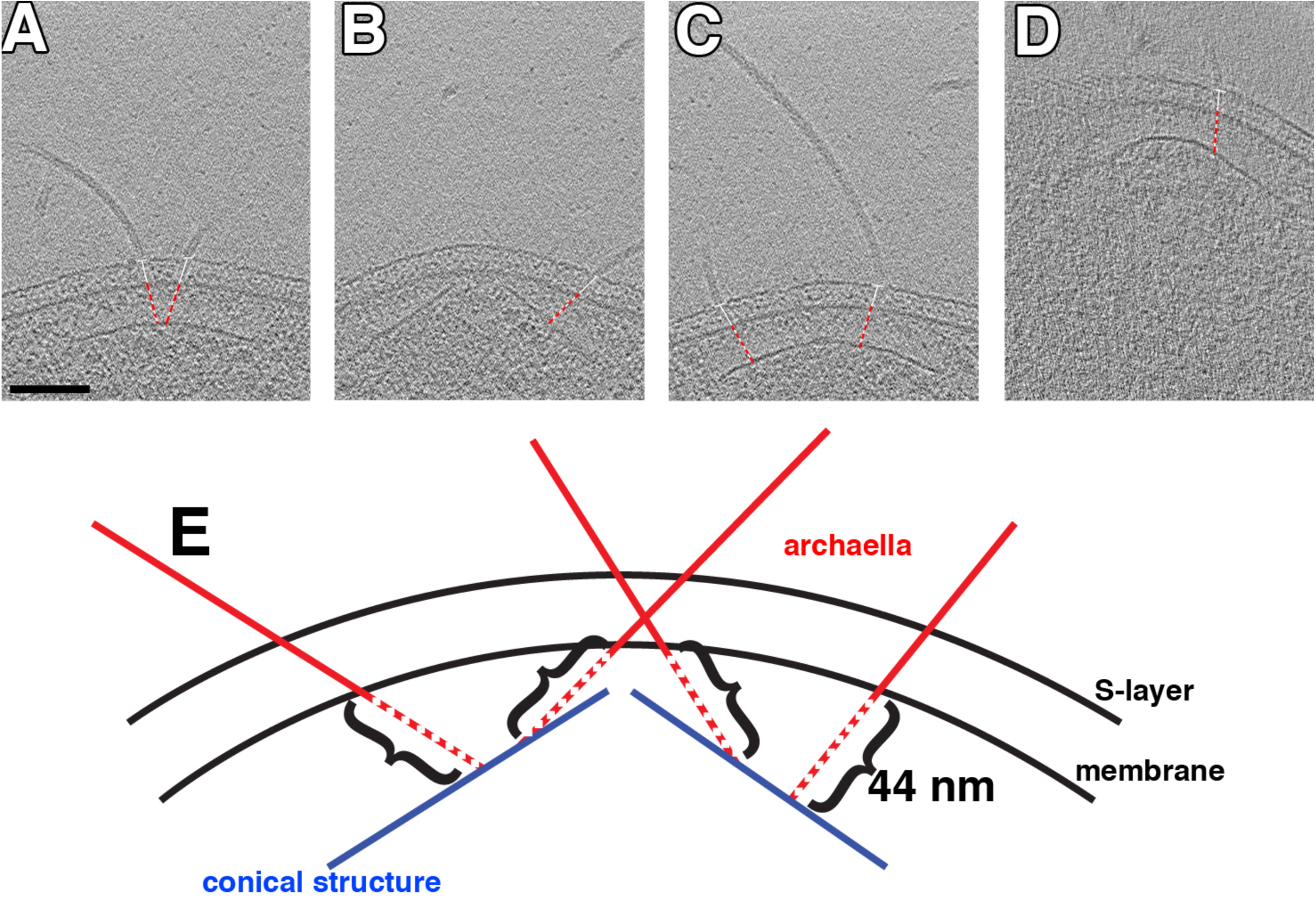
Archaellum orientation with respect to the cell envelope. **(A-D)** show tomographic slices through side views of cones. White lines show the angle of the archaellum with respect to the surface layer, and red dashed lines show the conserved distance from the archaellum at the membrane to the cone. Schematic in **(E)** depicts the 44 nm distance from the cone to the basal body in the membrane for archaella at different radial positions along the cone. Since different radial positions on the cone are located at different distances from the membrane (shorter at the tip and longer at the base), this results in a range of archaellar orientations in the cell envelope. Scale bar 100 nm.

To determine the structure of the archaellar basal body, we calculated a subtomogram average (Figure 4). 30 particles were used, and an axial two-fold symmetry was applied. The resulting average revealed several layers of density extending into the cytoplasm. Immediately adjacent to the membrane-embedded density was a ring-like structure (L1 in Figure 4A). Below the ring was a disk of similar diameter (L2), followed by a larger diameter component (L3) and finally, at a greater distance, a less well-defined density. This density was 44 nm away from the membrane, corresponding to the cone. Consistent with our observation that archaella exhibited various orientations with respect to the S-layer, we did not observe a strong density corresponding to the S-layer in the average. As seen in individual particles, the component in L3 does not appear to be a ring, but rather comprises distinct legs, seen on one or both sides, that appear symmetric in the average (Figure EV4). Similarly, the density of the cone is more prominent in individual particles; different angles of the structure in different particles wash out in the average (Figure EV4).

**Figure 4.**
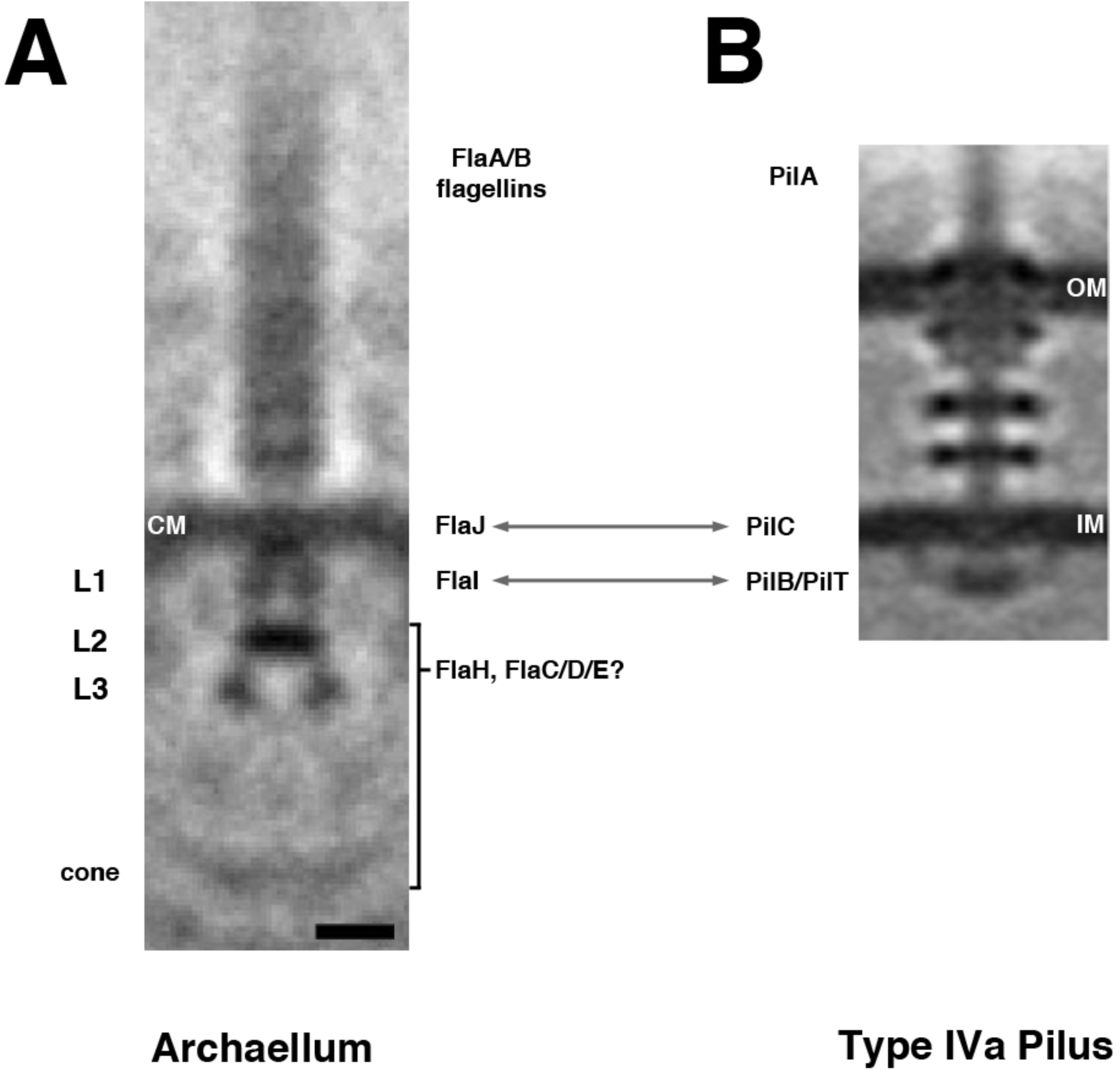
Structure of the *T. kodakaraensis*archaellum. **(A)** A sub-tomogram average of thearchaellum reveals structural features, including four layers of density in the cytoplasm (L1-L3, cone). CM, cytoplasmic membrane. The speculated identity of densities in the archaellum is proposed: archaellum fiber = FlaA/B flagellins; integral membrane density = FlaJ; L1 = FlaI; L2/L3/cone = FlaH/FlaC/D/E. **(B)** For comparison, a subtomogram average of the type IVa pilus machine from *Myxococcus xanthus* is shown (adapted with permission from [8]). Arrows indicate components with recognized homology. OM, outer membrane; IM, inner membrane. Scale bar 10 nm.

## DISCUSSION

### Structure of the basal body of the T. kodakaraensis archaellum

Here we describe the structure of the archaellar motor in *T.kodakaraensis* (Figure 4). We think it is almost certain that density L1 in the *T. kodakaraensis* basal body corresponds to the ATPase, FlaI, since its size and shape match those of the homologous ATPases in the T4P and all three interact directly with integral membrane proteins. More specifically, FlaI shares domain homology with the assembly/disassembly ATPases, PilB and PilT, of the bacterial T4P. The size of L1 is comparable to that of the PilB/PilT ring in the bacterial T4P (Figure 4B), consistent with their conserved hexameric oligomerization [7] and similar sizes of the protein monomers (540 amino acids for FlaI and 566 for PilB). FlaI is predicted to interact directly with the polytopic integral membrane protein FlaJ [6, 7]. FlaJ shares sequence homology with the ATPase-interacting inner membrane protein PilC of the T4P [18]. The relative locations of these components are therefore predicted to be the same in the basal bodies of the archaellum and the bacterial T4P (Figure 4) [8, 19], and the size and shape and position of density L1 seen here support that expectation, and the corollary that these two systems likely share a similar assembly mechanism.

Here we describe the structure of the archaellar motor in *T.kodakaraensis* (Figure 4). We think it is almost certain that density L1 in the *T. kodakaraensis* basal body corresponds to the ATPase, FlaI, since its size and shape match those of the homologous ATPases in the T4P and all three interact directly with integral membrane proteins. More specifically, FlaI shares domain homology with the assembly/disassembly ATPases, PilB and PilT, of the bacterial T4P. The size of L1 is comparable to that of the PilB/PilT ring in the bacterial T4P (Figure 4B), consistent with their conserved hexameric oligomerization [7] and similar sizes of the protein monomers (540 amino acids for FlaI and 566 for PilB). FlaI is predicted to interact directly with the polytopic integral membrane protein FlaJ [6, 7]. FlaJ shares sequence homology with the ATPase-interacting inner membrane protein PilC of the T4P [18]. The relative locations of these components are therefore predicted to be the same in the basal bodies of the archaellum and the bacterial T4P (Figure 4) [8, 19], and the size and shape and position of density L1 seen here support that expectation, and the corollary that these two systems likely share a similar assembly mechanism.

The identities of the proteins making up L2, L3, and the cone remain unclear. In the T4P, no structures were observed in the cytoplasm below the ATPase [8, 19, 20]. In Crenarchaeota, only one accessory component is not membrane-bound (FlaH). In Euryarchaeota like *T.kodakaraensis*, however, additional soluble proteins, FlaC/D/E, are thought to be components of the archaellum that receive switching signals from the chemotaxis machinery [21]. All of these proteins, and potentially others, are candidates for the densities we observed. It will therefore be of great interest to obtain a structure of the crenarchaeal basal body, which lacks FlaC/D/E, for comparison, and/or to dissect the *T. kodakaraensis* basal body structure through analysis of deletion mutants.

### Conical structures anchor T. kodakaraensis archaella

We observed that *T. kodakaraensis* archaella associate with a large conical structure in the cytoplasm. In a few cases, we observed direct connections between archaella and cone. The fact that we did not see such a connection for every archaellum may simply reflect variations in image clarity and orientation of the structures between cells in different cryotomograms. The conserved distance from the cone to the archaellar basal body in the membrane suggests a rigid interaction. It is an interesting question how archaella are attached to the cone. We did not observe strong densities connecting L3 and the cone in the averaged basal body structure, but in individual particles we observed heterogeneity. Also, the resolution of the average may be too low to detect such connections. If, for example, the links are thin (such as coiled-coils), they would not be resolved; similar coiled-coil linkages in the bacterial flagellar motor between FlaH and the C-ring were not resolved even in higher-resolution subtomogram averages [22].

We propose that the *T. kodakaraensis* cone anchors the archaellar basal body in part to provide leverage for rotation. In the bacterial T4P, the ATPase is clamped by extensive interactions up through the cell envelope that anchor it to the peptidoglycan cell wall [8] (Figure 5). Signals governing disassembly are thought to be processed by sensory elements in the periplasm [8]. In the absence of a peptidoglycan cell wall and outer membrane, the *T. kodakaraensis* archaellum appears to turn the system upside down, with components stacking nearly 50 nm into the cytoplasm to anchor onto a large cone (Figure 5). Signals governing rotation and direction are likely integrated by sensory components in the cytoplasm.

**Figure 5.**
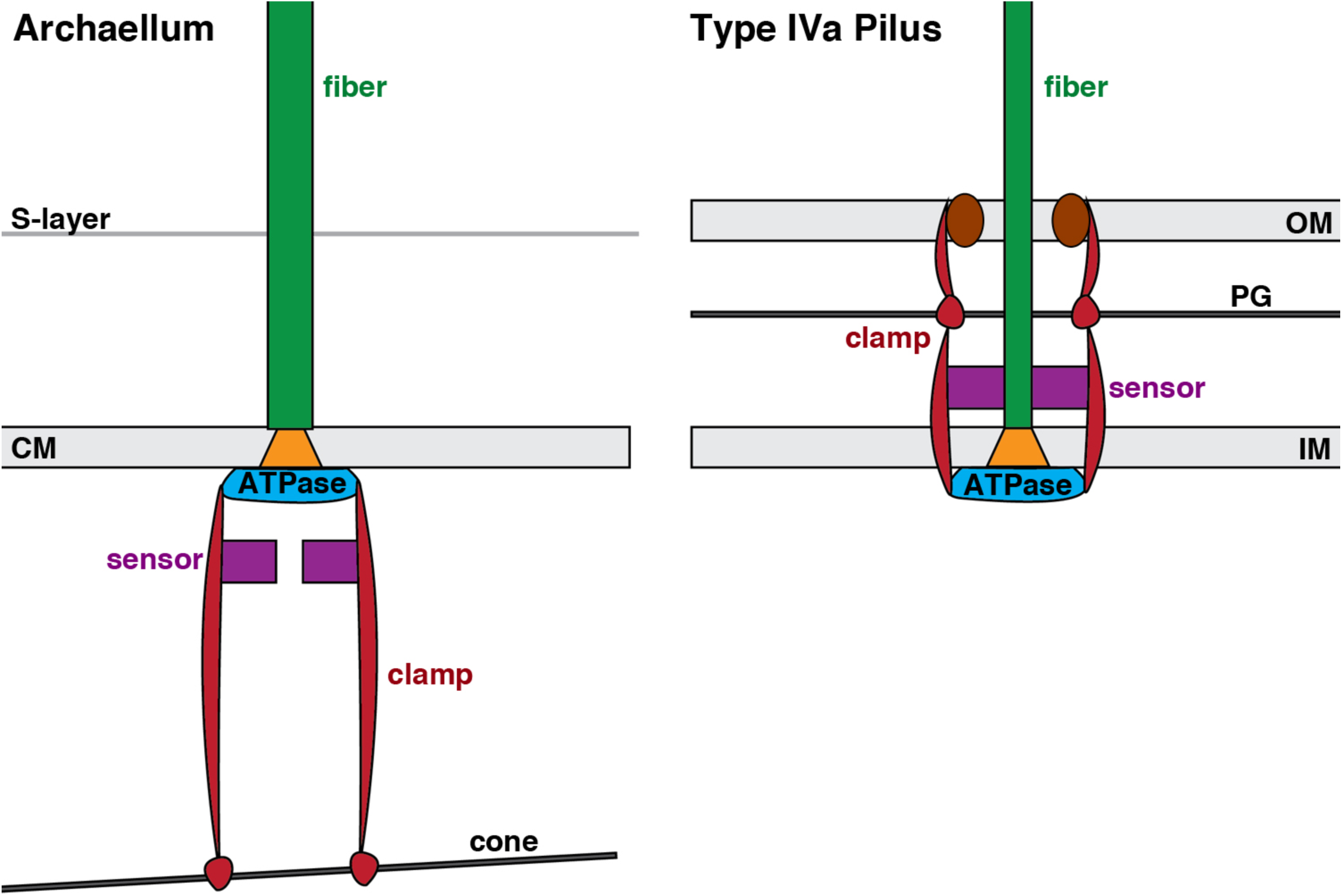
Schematic comparing organization of the related archaellum and type IVa pilus basal bodies. In the bacterial T4P **(right)**, an integrated system of components spanning theouter and inner membranes (OM, IM) uses the peptidoglycan cell wall (PG) to brace the ATPase, allowing rotation of PilC (orange) in the membrane to assemble the pilus fiber. In the *T.kodakaraensis* archaellum**(left)**, our results suggest that an integrated system of componentsextends from the single membrane (CM) inward to a large conical structure in the cytoplasm to similarly brace the ATPase. Sensory components (purple) are proposed to be located in the periplasm for the T4P and the cytoplasm for the archaellum.

Similar leveraging structures may exist in other Archaea. More than twenty years ago, it was observed that when archaella are dissociated from lophotrichous *H. salinarum* cells by detergent, the bundles remain intact, connected to a large (~500 nm diameter) structure [11]. A similar structure was also observed below the cell membrane in cell ghosts [12]. More recently, a spherical structure was observed anchoring Iho670 fibers, T4P-like filaments in *Ignicoccus hospitalis*. This structure is thought to be located in the cytoplasm of the cell and contains a central ring of similar dimensions to the one observed here [23]. It is possible that either or both of these structures are related to the *T. kodakaraensis* cone. Large cytoplasmic structures have not been described in other motile archaeal species to date, however, so it will be interesting to determine how archaella may be anchored in those systems.

It will also be of great interest to identify the proteins that form the *T. kodakaraensis* cone and associated ring. These subunits must be capable of interacting both circumferentially around the cone as well as radially with subunits making up the next (larger or smaller diameter) ring. While it is possible that the conical structure is an assembly of stacked rings, we think it more likely that the subunits assemble into a filament spiral, similar to what has been proposed for ESCRT-III polymers [24, 25]. Interestingly, an architecturally similar spiral has been observed in the basal body of the bacterial flagellar motor: in *Wolinella succinogenes*, an Archimedian spiral forms a bushing for the motor in the periplasm, allowing the flagellum to rotate in the cell wall. This spiral is formed by protein subunits interacting both circumferentially and laterally through nonspecific interactions [26]. While that spiral takes the form of a disk, similar protein interactions may give rise to a cone in *T. kodakaraensis*.

### T. kodakaraensis *cones are potential polar organizing structures*

In addition to a potential role in rigidly anchoring the basal body of the archaellum, the *T. kodakaraensis* cone may function to gather the archaellar bundle to maximize efficiency, either by concentrating molecules for assembly or signaling, or by concentrating force at one point on the coccoid cell for directional swimming. The cone’s structure may also help distribute the force from archaellar rotation to the larger bulk of the cell’s contents. This might be more efficient in a pushing than a pulling mode; swimming speed in another euryarchaeon, *H. salinarum*, was found to be approximately twice as fast when the archaella push as when they pull the cell body [5]. The structurally-similar spiral basal disk in the bacterial flagellar motor of *W. succinogenes* was suggested to play a role in dispersing lateral forces created by flagellar rotation [26].

Our results suggest a further role for the cone in breaking the symmetry of the coccoid cell. In many rod-shaped bacterial cells, proteins and other macromolecules are specifically localized to the cell pole for various purposes ranging from cell motility and adhesion to differentiation and division [27]. One well-studied example of this polar organization occurs in *Caulobacter crescentus*, where the oligomeric protein PopZ defines an asymmetric pole, localizing many cytoplasmic proteins and tethering the chromosomal centromere to facilitate division [28– 30]. In *Vibrio cholerae*, the HubP protein organizes the polar localization of the chromosomal origin, chemotaxis machinery, and flagella [31]. Perhaps the *T. kodakaraensis* cone similarly defines a pole in the spherical cells, anchoring the chemotaxis and motility machinery. An intriguing feature observed in our cryotomograms is the cone-associated REZ. In bacterial cells, such REZs are commonly interpreted to be the nucleoid [32, 33]. Supporting this assignment, we observed bundles of filaments (most extensive in lysed cells) associated with the REZ (Figure 2). Such filaments are reminiscent of nucleoprotein filaments formed by various bacterial DNA-binding proteins in stress conditions [34–36].

A spatial organizer analogous to PopZ may be especially important for a polyploid species like *T. kodakaraensis* (chromosome copy number varies depending on growth phase, from 7 to 19 copies [37]). Fluorescence imaging suggests that the nucleoid is relatively compact in log phase growth, and nucleoids appear to separate before the cells are deeply constricted [38]. Perhaps the cones segregate attached structures, including the archaella and possibly chromosomes. This function is consistent with the duplicated cone structure we observed in one cell (Figure EV2), which could represent an intermediate after replication and prior to segregation, or may simply represent an aberrant structure. Further studies imaging cells throughout the cell cycle could shed light on whether, and how, cones function to coordinate archaellar and chromosomal segregation.

Understanding the prevalence of this structure among Euryarchaeota and across different archaeal kingdoms may illuminate its function. If it is restricted to lophotrichous species, it may simply be an anchoring mechanism for the archaella in the absence of a peptidoglycan cell wall. In that case, monotrichous or peritrichous species may exhibit a less extensive plate underneath the basal bodies of individual archaella. If it is more widely found in coccoid, and/or highly polyploid, cells, it may serve an added role in polar specification.

## METHODS

### Growth

*Thermococcus kodakaraensis* strain KOD1 [JCM 12380] was grown anaerobically in MA-YT medium supplemented with elemental sulfur as previously described [14, 39].

### Electron cryotomography and image analysis

Samples of cell cultures in growth media were mixed with bovine serum albumin-treated colloidal gold fiducial markers (Sigma) and applied to Quantifoil R2/2 200 copper EM girds (Quantifoil Micro Tools). After blotting excess liquid, grids were plunge-frozen in a mixture of liquid ethane and propane [40], and subsequently kept at liquid nitrogen temperature. Images were acquired using either an FEI Polara G2 or Titan Krios 300 keV transmission electron microscope (FEI Company) equipped with a field emission gun, image corrector for lens aberration, energy filter (Gatan), and K2 Summit direct electron detector (Gatan). Cumulative electron dose was 160 e^-^/Å^2^ or less for each tilt-series. Tilt-series were acquired using UCSF Tomography software [41]. Images were contrast transfer function corrected, aligned, and reconstructed by weighted back projection with the IMOD software package [42]. SIRT reconstructions were calculated with TOMO3D [43], subtomogram averages generated using PEET [44], and segmentations generated with Amira software (FEI Company).

## ACCESSION CODES

The subtomogram average of the *T. kodakaraensis* archaellar basal body was deposited into the Electron Microscopy Data Bank (entry number EMD-8603).

## ACKNOWLEDGMENTS

This work was supported by the NIH (grants R01GM101425 and R01AI127401 to G.J.J.) and the UCLA-DOE Institute (grant DE-FC03-02ER26342 to R.P.G.).

---

**Figure EV1.**
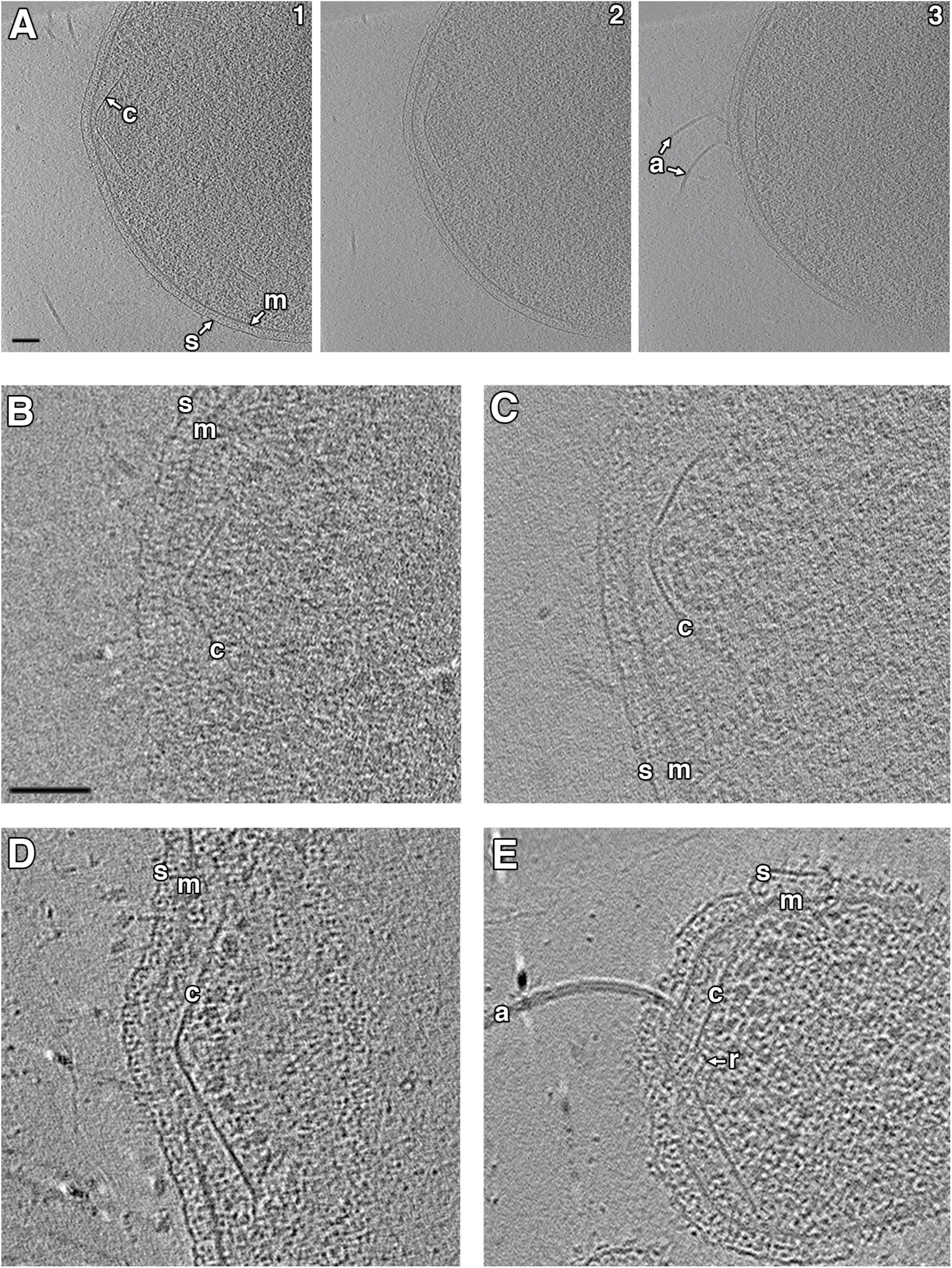
Additional examples of conical structures in*T. kodakaraensis*cells. **(A)** shows sequential tomographic slices (1–3) at different heights through a side view of the cone. Additional examples of cones in intact **(B-C)** and lysed **(D-E)** cells are shown below. s, S-layer; m, membrane; c, conical structure; a, archaella; r, ring. Scale bars 100 nm.

**Figure EV2.**
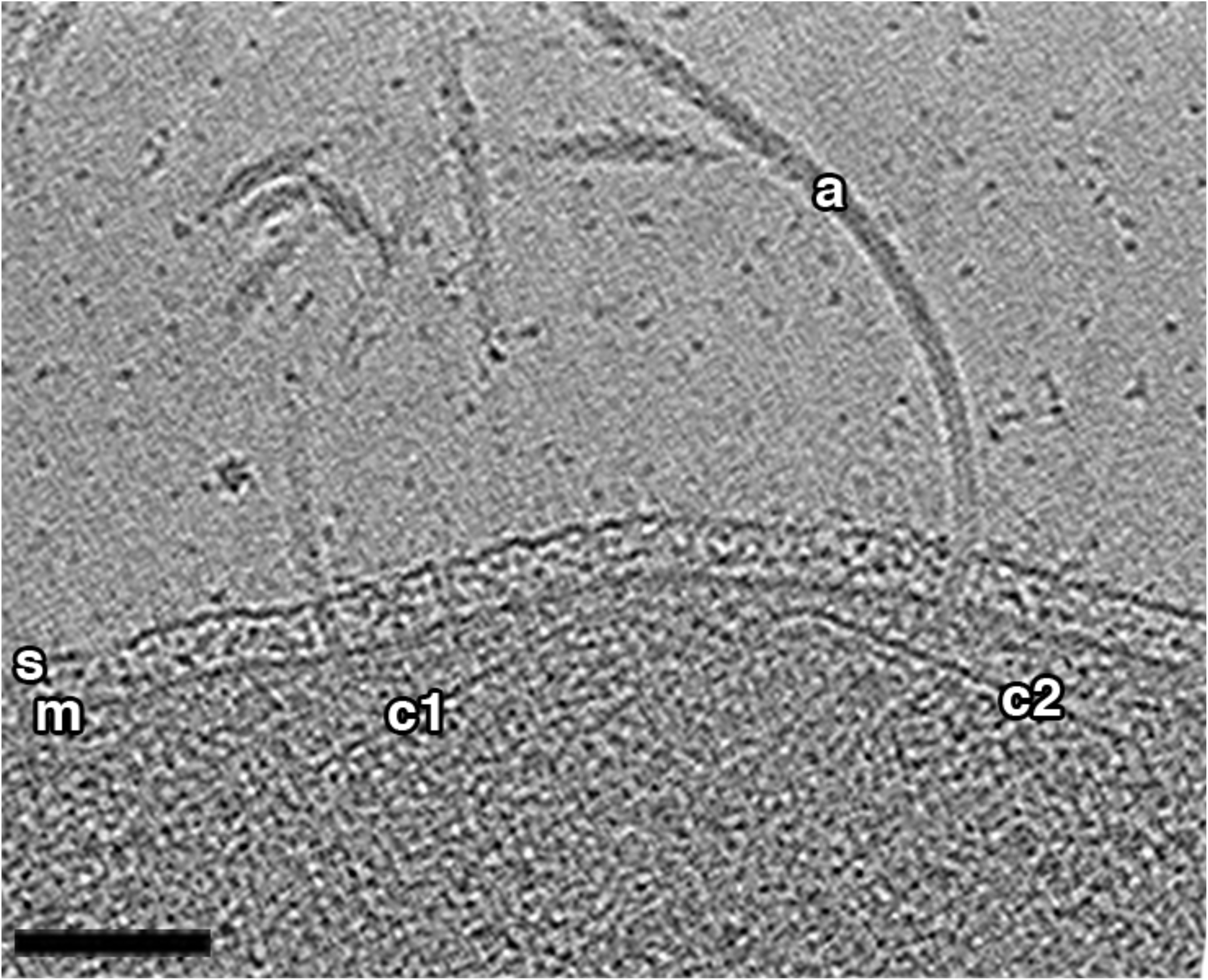
Double cone structure observed in *T. kodakaraensis*. A tomographic slice through a side view shows two associated conical structures (c1 and c2), both associated with archaella (a). Scale bar 100 nm.

**Figure EV3.**
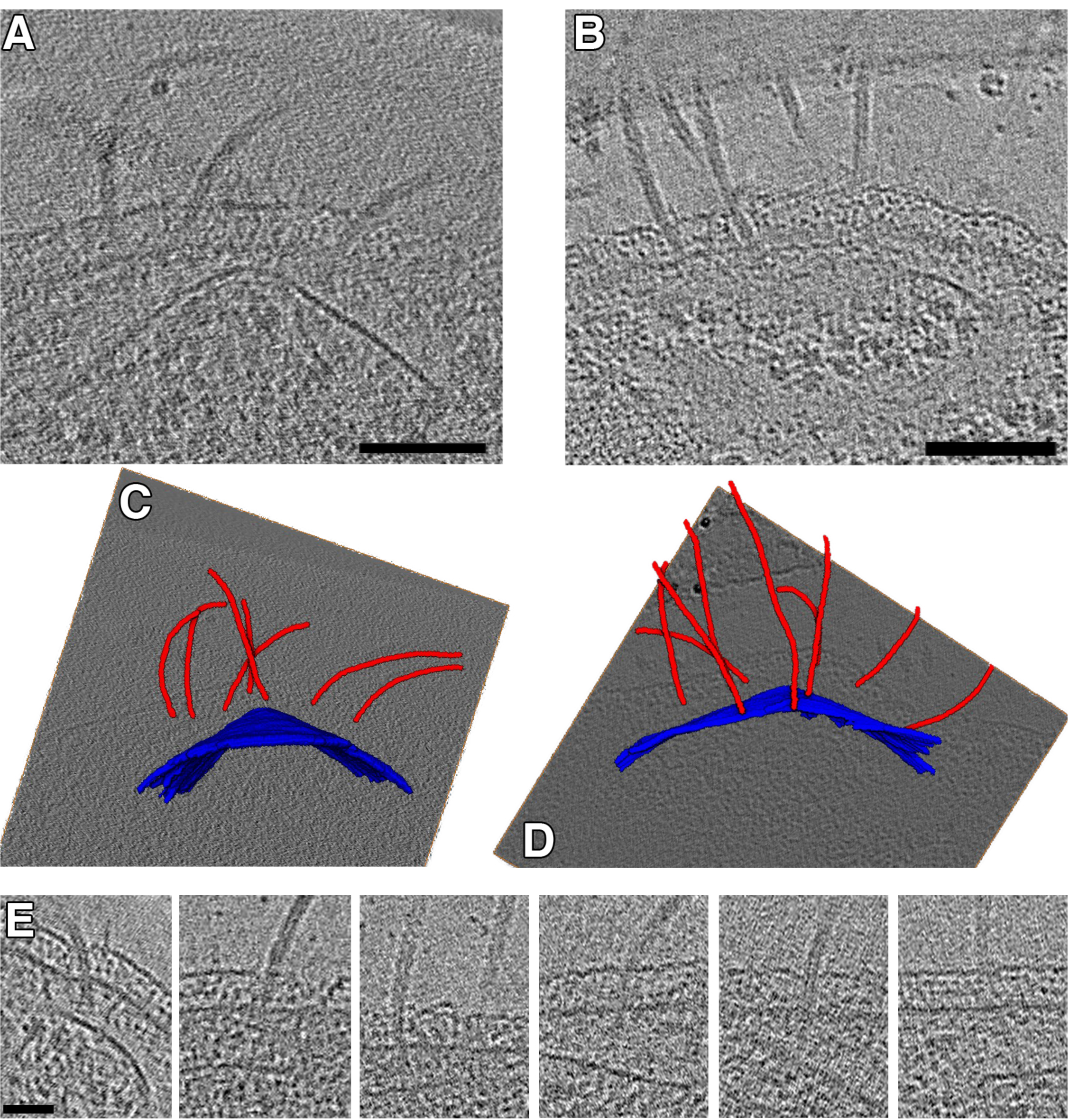
Association of cones with the archaellar bundle. **(A)** and **(B)** show tomographic slices through two cells, highlighting the association between the cone and the archaella. **(C)** and show 3D segmentations of the cells in **(A)** and **(B)**, respectively, with cones in blue and archaella in red, embedded in tomographic slices. **(E)** Tomographic slices of individual archaella show the varying orientations of archaella with respect to the cell envelope, as well as apparent connections to the cone. Scale bars 100 nm in (A) and (B), 50 nm in (E); segmentations not to scale.

**Figure EV4.**
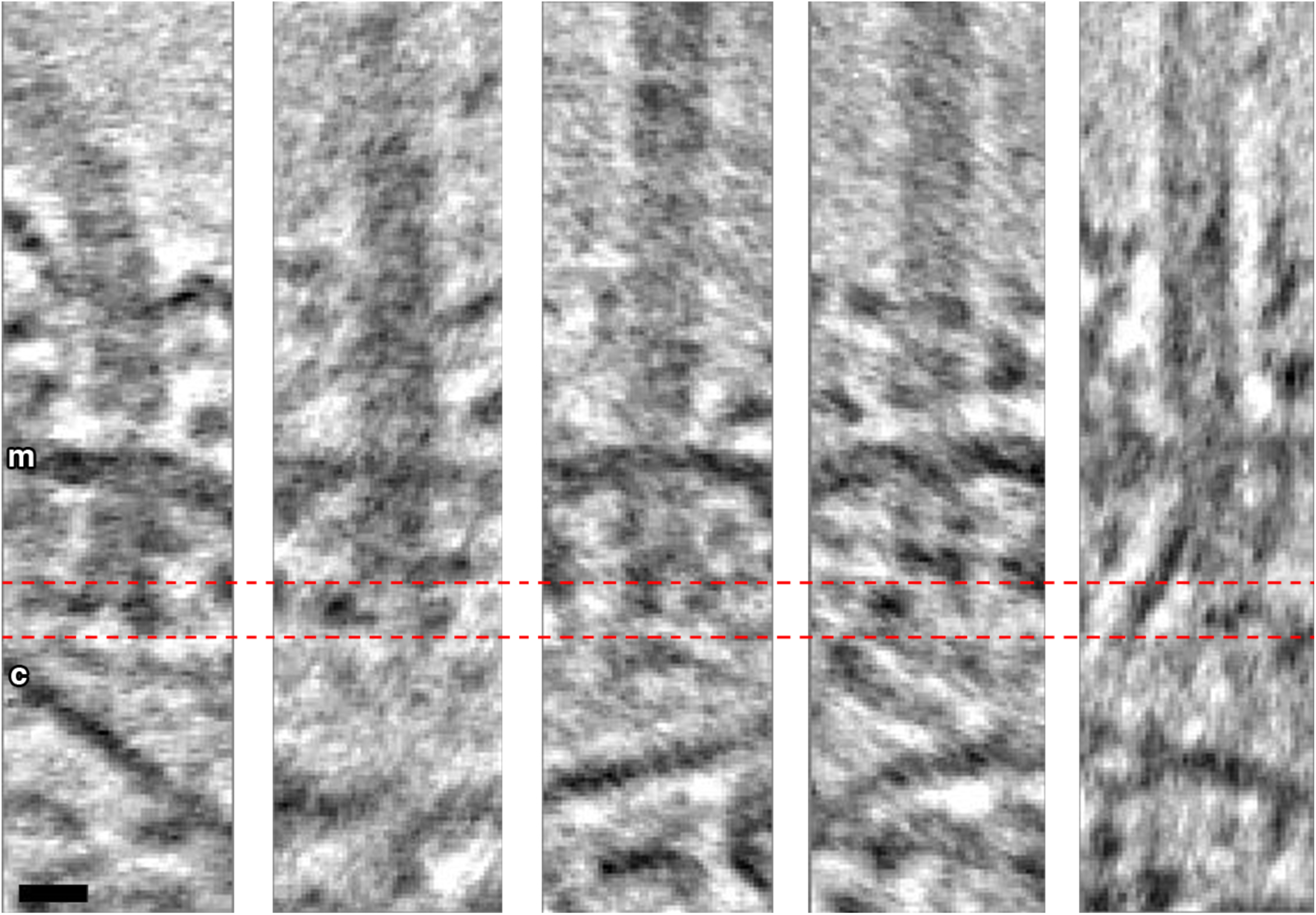
Individual particles from the subtomogram average show heterogeneity in the L3 density and angle of cone density. The L3 density appears as either two dots of similar(first two panels) or different intensity (third panel), a single dot (fourth panel), or a dot and an extended line (fifth panel). Scale bar 10 nm.

